# Symbiotic Bacterial Partners for Chemical Defense are Vertically Transmitted in a Marine Sponge

**DOI:** 10.1101/2024.12.29.630706

**Authors:** Masaki Fujita, Yuji Ise, Masashi Fukuoka, Nobutada Kimura, Akihiro Ninomiya, Kazutoshi Yoshitake, Shigeki Matsunaga, Ryuichi Sakai, Kentaro Takada

**Affiliations:** Graduate School of Fisheries Sciences, Hokkaido University; Hakodate, Hokkaido 041-8611, Japan; Kuroshio Biological Research Foundation; Otsuki-cho, Kochi 788-0333, Japan; Graduate School of Science, Nagoya University; Toba, Mie 517-0001, Japan; Bioproduction Research Institute, The National Institute of Advanced Industrial Science and Technology; Sapporo, Hokkaido 062-8517, Japan; Graduate School of Agricultural and Life Sciences, The University of Tokyo; Bunkyo-ku, Tokyo 113-8657, Japan; School of Marine Biosciences, Kitasato University; Sagamihara, Kanagawa 252-0373, Japan

## Abstract

Marine sponges host diverse symbiotic bacteria, some of which are essential for chemical defense. However, the mechanisms governing the inheritance of these defensive symbionts remain elusive. Here we report the first case of a transmission mechanism for symbionts producing cytotoxins in the sponge *Mycale nullarosette*, for which we established the full life-cycle cultivation method. The identification of *Candidatus* Synechomycale tutelaris as the cytotoxins producer enabled us to unveil their vertical transmission to offspring via bacteriocyte-like structures from the embryonic stage. This study demonstrates that sponges have evolved a sophisticated system to ensure the inheritance of significant bacterial partners crucial for host defense, which is likely one of the reasons that sponges have survived throughout the long history of multicellular organisms.

## Introduction

Marine sponges, among the earliest branching metazoans, are one of the most primitive multicellular organisms on Earth (*1*). These organisms play an essential role in the nutrient cycling within benthic communities, primarily through their efficient filter-feeding of dissolving organic materials and microorganisms (*2, 3*). Many sponges harbor diverse and abundant microbial communities, which constitute up to 40% of the sponge biomass (*4*), making them ideal models for studying animal-microbe symbiosis.

From a chemical perspective, marine sponges are prolific sources of bioactive compounds with significant potential for drug development (*5*). Many of these metabolites are attributed to the symbiotic microorganisms within sponges, due to their structural similarity to compounds produced by terrestrial bacteria (*6, 7*). For instance, symbionts of the candidate genus ‘Entotheonella’ have been genetically identified as the producers of various bioactive natural products in sponges *Theonella swinhoei* and *Discodermia calyx* (*8*–*11*). Similarly, in *Mycale hentscheli*, several bacteria, such as *Candidatus* Patea custodiens, *Ca*. Entomycale ignis, and *Ca*. Caria hoplite, have been identified as responsible for the production of cytotoxic metabolites (*12, 13*). In the case of *Haliclona* sp., symbiotic *Ca*. Endohaliclona renieramucinifaciens was identified as a producer of renieramycin (*14*). These biosynthetic studies suggest that specific symbionts contribute to the chemical defense of the host sponges by producing toxic substances. However, the mechanisms by which sponges discriminate between bacteria for feeding and those for chemical defense remain largely unexplored. Numerous studies have suggested that sponge-associated bacteria are acquired through a combination of vertical and horizontal microbial transmission (*4, 15, 16*).

To investigate the transmission mechanisms of symbionts involved in chemical defense, we focused on the marine sponge *Mycale nullarosette* (**Fig. 1A**). This species contains trisoxazole-type macrolides, mycalolides (**Fig. 1G, Fig. S1**), which are widespread among several sponge species and nudibranchs and are known for their role in chemical defense, including predation inhibition, cytotoxicity, and antifungal activity (*17, 18*). *M. nullarosette* is particularly well-suited for such studies due to its annual lifespan, ovoviviparous reproduction, and the availability of a full life-cycle cultivation method established by our groups (**Fig. 1B-F, Fig. S2**).

**Fig. 1.**
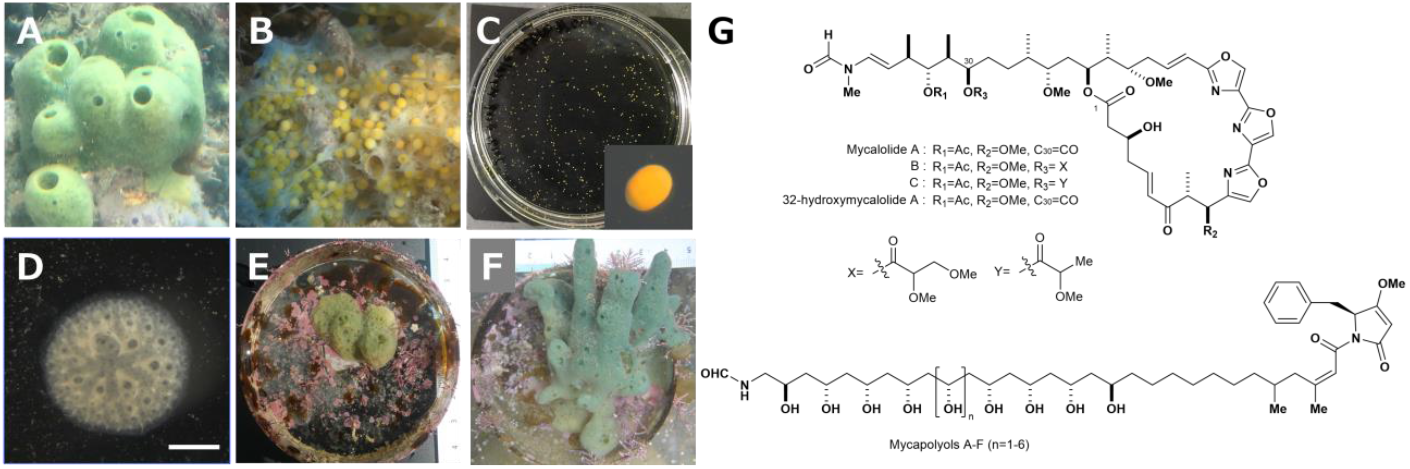
Marine sponge *Mycale nullarosette* and its cytotoxic metabolites, mycalolides and mycapolyols. (A) Mature individuals prior to propagation. (B) Embryos inside of the mature sponge (yellow particles about 0.5 mm diameter). (C) Swimming larvae released from the mature sponge, about 1.0 mm diameter. (D) Juvenile sponges one week after settlement on a Petri dish (scale bar: 1.0 mm), (E) six months, (F) eleven months after settlement, respectively. (G) Chemical structures of mycalolides and mycapolyols.

In this study, we initially identified the biosynthetic gene cluster (BGC) for mycalolides and determined the taxonomy of the producing bacterium using metagenomic analyses. We also examined the presence of the bacterium throughout the sponge’s life stages encompassing adults, embryos, larvae, and juveniles. Finally, microscopic observations, including fluorescence *in situ* hybridization (FISH) and transmission electron microscopy (TEM), clarified the transmission and localization mechanisms of the mycalolides-producing symbiont within *Mycale nullarosette*.

## Results

### Identification of the BGC for mycalolides and mycapolyols

To identify the BGC responsible for mycalolides, we performed whole metagenome sequencing of the bacterial fraction from *M. nullarosette* using the Illumina platform. This analysis yielded approximately 5,000 contigs totaling 40.6 Mb, among which ten contigs contained PKS-NRPS related genes (**Table S1**). Based on the PKS-NRPS domain architecture of each contig, the predicted products of five contigs were in good agreement with the core structure of mycalolides, notably with the characteristic trisoxazole moiety derived from three consecutive serine residues. Binning analysis, based on the coverage depth and GC content, revealed that these five contigs were identified as part of a single cluster, designated the *myl* cluster (**Fig. S3**). PCR gap closure revealed that the *myl* cluster comprises five genes (*mylA*-*E*), spanning 105 kb (**Fig. 2A**). The domain arrangement of the *myl* genes matched well with mycalolides biosynthesis (**Fig. S4A, Table S2**).

**Fig. 2.**
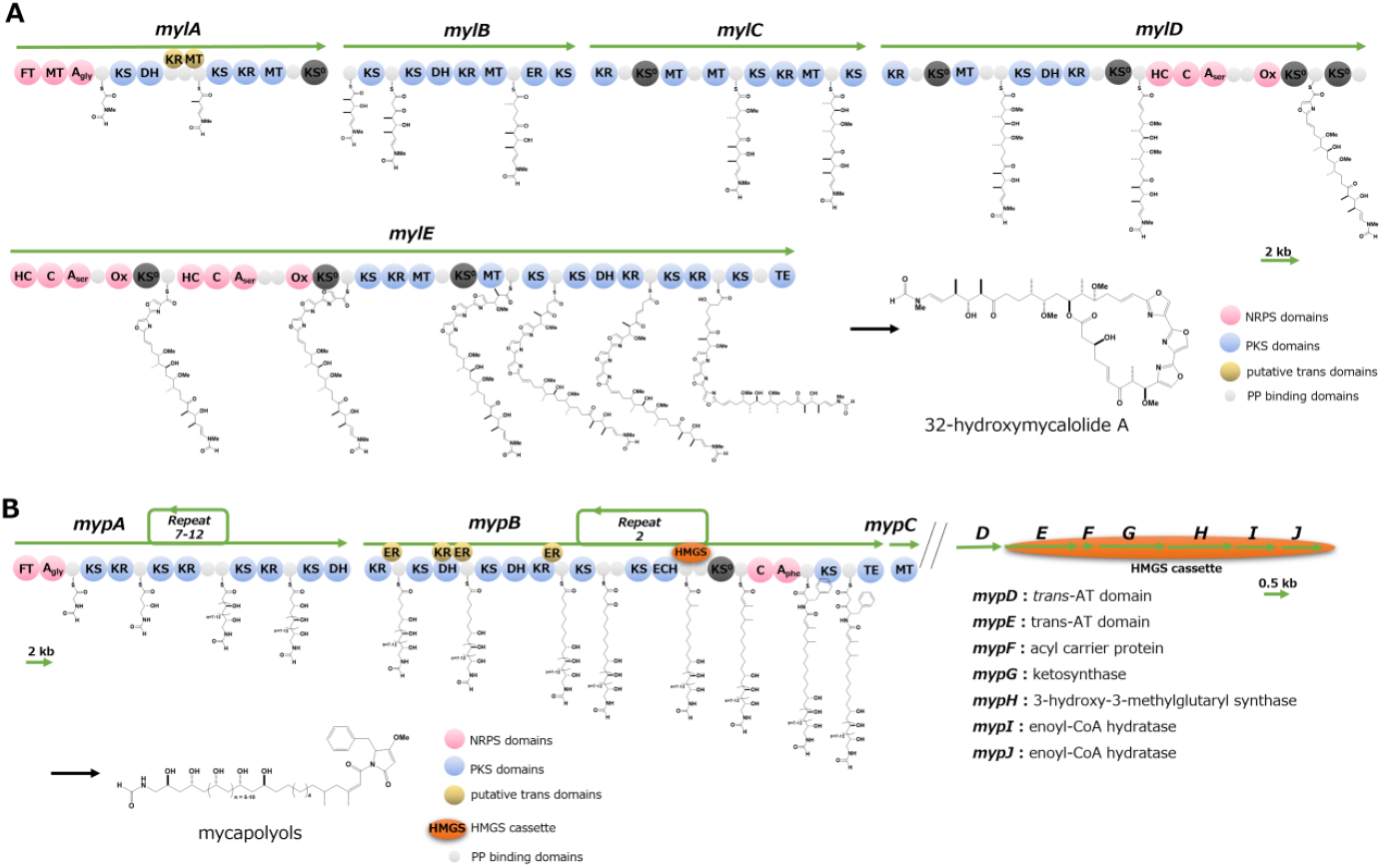
Overview of the putative biosynthetic gene clusters for mycalolides and mycapolyols. (A) Domain architecture of the *myl* cluster and the predicted biosynthetic scheme for 32-hydroxymycalolide A, an initial biosynthetic product of the mycalolide series. (B) Domain architecture of the *myp* cluster, and the predicted biosynthetic scheme for mycapolyols.

MylA starts with the adenylation domain (A_gly_) to incorporate glycine, followed by *N*-formyltransferase (FT) and *N*-methyltransferase (MT) domains that form the *N*-formyl-*N*-methylglycine moiety, a terminal structure of mycalolides. Additionally, MylD and E contain three successive A domains (A_ser_), which are accompanied by oxidation (Ox) and heterocyclase (HC) domains to transform serine residues into oxazoles (*19*). Among the 22 ketosynthase (KS) domains identified, nine appear to be non-elongating KS^0^ domains (**Fig. 2A, Table S3**), consistent with the structural features of mycalolides. Interestingly, MylA lacks one keto-reductase (KR) and one MT domain, a deficiency similar to that observed in the BGCs of luminaolide (*lum* cluster) and tolytoxin (*tto* cluster) (*9*), cyanobacterial products with side chains nearly identical to that of mycalolides (**Fig. S5**). For the *lum* and *tto* clusters, it is speculated that *trans*-type KS and MT enzymes, encoded outside of the BGC, compensate for this deficiency.

Beyond the *myl* cluster, we identified two additional contigs encoding PKS-NRPS in the metagenome. These genes corresponded to the biosynthesis of mycapolyols (**Fig. 1G**), unique PKS-NRPS metabolites previously isolated from the other species *Mycale izuensis* (*20*). The predicted structure of mycapolyols included a continuous 1,3-polyol system starting with *N*-formylglycine and a polyene segment terminating in a γ-lactam formed from a phenylalanine residue (**Fig. S4B**). Mycapolyols include analogues with nine to fourteen successive secondary hydroxy groups, suggesting biosynthesis by an iterative-type PKS. The relative configurations of 1,3-diol moieties in mycapolyols have been determined to be *syn*, except at both ends (*20*) The stereochemistry of the three KR domains is predicted to be *R, S*, and *R*, respectively, due to the presence or absence of Asp residue in the LDD motif (*21*). This indicates that the first and third KR domains contribute to the formation of hydroxy groups at both ends, while the inner hydroxy groups are formed by the second KR domain in an iterative mode. Additionally, one contig contained a 3-hydroxy-3-methylglutaryl synthase (HMGS) cassette involved in constructing β-branched polyketides. The gene arrangement, designated *myp A-J*, perfectly matched the structure of mycapolyols (**Fig. 2B**). The presence of ion peaks corresponding to mycapolyols A-D in *M. nullarosette* extract confirmed their biosynthesis and further supported the functional annotation of the *myp* cluster (**Fig. S6**).

### Whole genome sequence to identify the mycalolides producer

To identify the mycalolides producer, we employed long-read whole metagenome sequencing to determine its complete genome. High molecular weight DNA prepared from the bacterial suspension of *Mycale nullarosette* was sequenced using MinION, yielding a circular draft genome, which was subsequently refined with Illumina Hiseq reads to produce a complete 2.53 Mb genome sequence (accession: AP024111; **Fig. 3A**). The closest culturable species with complete genome data is *Coraliomargarita akajimensis* DSM45221, belonging to the phylum Verrucomicrobia with 87% identity on 16S rRNA, and an ANI value 64.4 (**Table S4**) (*22*). Phylogenetic analysis using SILVA 16S rRNA database suggests that the mycalolides producer represents a novel genus within the phylum Verrucomicrobia, specifically the class Opitutae, order Opitutales, and family Opitutaceae (**Fig. 3B**). We propose naming it *Candidatus* Synechomycale tutelaris strain Am1 (tutelaris, meaning ‘guardian’ in Latin). Another genome sequence of the mycalolides producer, derived from *M. nullarosette* collected at a different location, is named as *Ca*. S. tutelaris strain Sg1 (accession: AP024164), possessing a nearly identical sequence to that of strain Am1 (**Table S4**).

**Fig. 3.**
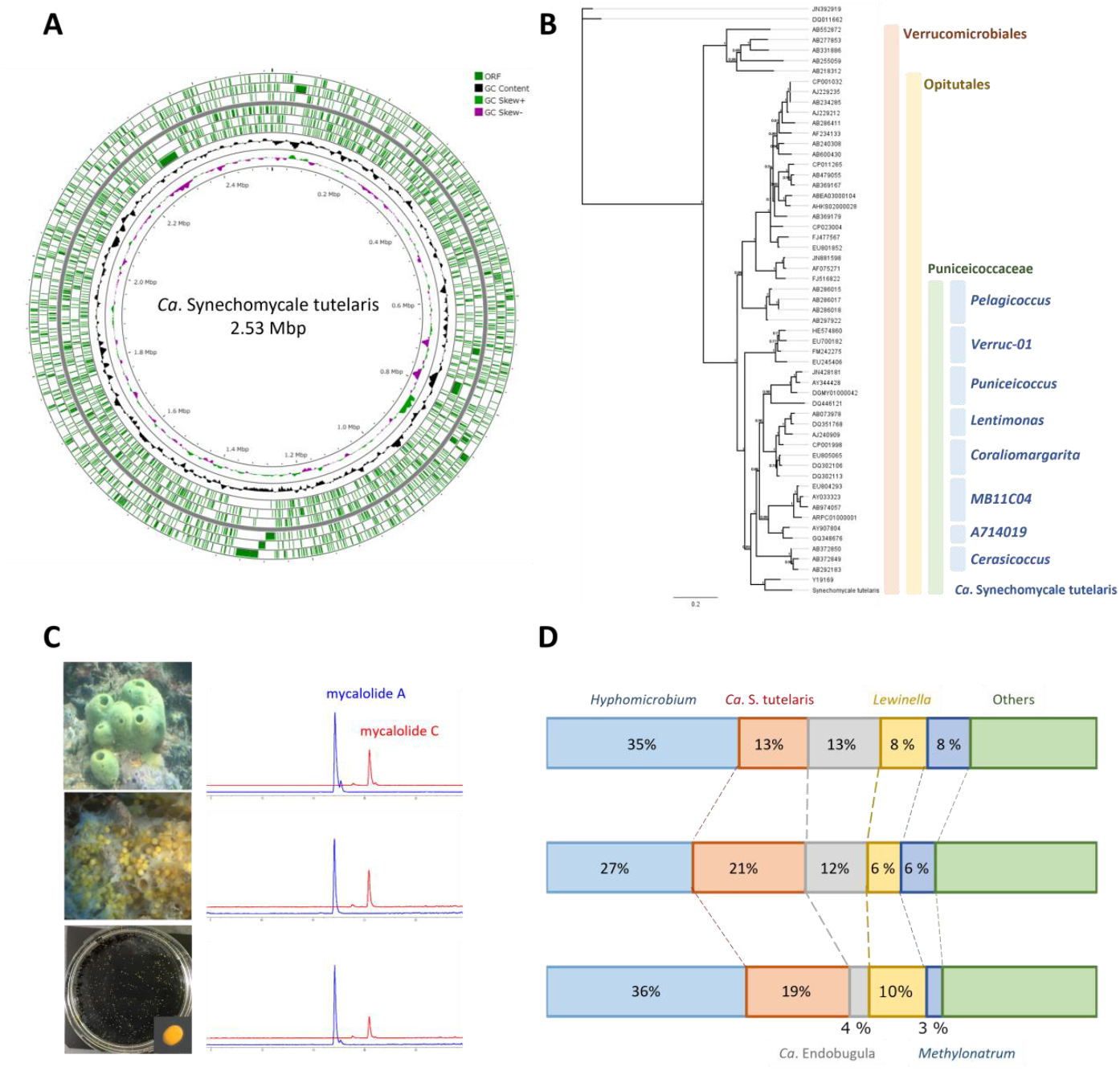
Sponge symbionts, *Ca*. S. tutelaris, and its distribution across life cycles. (**A**) Genome map of the mycalolides producer, *Ca*. S. tutelaris. (**B**) Phylogenetic tree of 16S rRNA sequences from bacteria within the class Verrucomicrobiales. (**C**) Chemical profile of mycalolides observed in adult sponge (top), embryos (middle), and larvae (bottom), (**D**) Bacterial flora in adult sponge (top), embryos (middle), and larvae (bottom).

The annotation of the ‘Synechomycale’ genome by Rapid Annotation using Subsystem Technology (RAST) (*23*) identified 2,987 coding sequences (CDSs, **Table S5**). Intriguingly, ‘Synechomycale’ lacks many essential genes involved in primary metabolic processes, such as carbohydrates, amino acids, lipids, and nucleotides, as well as other vital genes for respiration and stress response. Notably, only four CDSs related to ‘cell wall and capsule’ (compared to 32-81 CDSs in representative culturable Verrucomicrobium bacteria) and no CDS for ‘cell division and the cell cycle’ (11-29 CDSs in related species) was identified. In contrast, the genome contains 291 CDSs associated with mobile genetic elements, such as transposases and integrases, significantly more than the culturable Verrucomicrobia species (4-16 CDSs). These data suggest that many essential genes in ‘Synechomycale’ have become non-functional due to the insertions of mobile genetic elements, leading to formation of pseudogenes. This is a hallmark that the genome is undergoing a shrinking progress.

Despite the loss primary metabolic functions, ‘Synechomycale’ genome retains an enrichment of secondary metabolism genes, such as *myl* and *myp* clusters, accounting for 8.7% of the entire genome. Moreover, all PKS-NRPS genes found in the whole sponge metagenome are exclusive to the ‘Synechomycale’ genome (**Table S2**). These data imply that ‘Synechomycale’ plays a pivotal role in the chemical defense of its host, and the host provides a niche for its independent symbiont.

### Mode of transmission of *Ca*. S. tutelaris

The strong symbiotic relationship between ‘Synechomycale’ and *M. nullarosette* motivated us to investigate how the sponge acquires the symbionts. By adapting the aquaculture techniques used for the ascidian *Ciona intestinalis*, we have successfully cultivated *M. nullarosette* throughout its entire life cycle, obtaining specimens from adults, embryos, larvae, and second-generation juvenile sponges (**Fig. 1B-F, Fig. S2**). Larvae were settled on Petri dishes and cultivated in an indoor tank, an outdoor tank, and a shaded cage in the sea, yielding juvenile sponges. LC-MS analyses for the extracts from each life stage showed consistent ratios of mycalolides throughout the entire life cycle (**Fig. 3C, Fig. S7**). To confirm the presence of ‘Synechomycale’ across all life stages, we performed 16S rRNA amplicon sequencing on metagenomic DNA collected from each stage. ‘Synechomycale’ was consistently detected, alongside other major symbionts belonging to the genus *Hyphomicrobium, Ca*. Endobugula, *Methylonatrum*, and *Lewinella* (**Fig. 3D, Table S7**). In contrast, targeted amplification using primer sets designed for the *myl* cluster failed to detect ‘Synechomycale’ in surrounding environmental DNA, or a nearby unidentified sponge (**Table S8**), suggesting that ‘Synechomycale’ is neither free-living nor an opportunistic symbiont (**Fig. S8**). Shotgun metagenomics further confirmed the presence of complete *myl* clusters in larvae. These data show that ‘Synechomycale’ constructs a highly specific symbiotic relationship with *M. nullarosette*, and establishes a vertical transmission pathway that begins as early as the embryonic stage.

### Localization of *Ca*. S. tutelaris in *M. nullarosette*

Generally, symbiotic bacteria in sponge hosts have been reported to reside either in the extracellular matrix or within bacteriocytes (*4, 24*). However, the specific localization of ‘Synechomycale’ at the early developmental stage remains unknown. To reveal bacterial localization in embryo and larvae, we conducted fluorescence *in situ* hybridization (FISH) targeting its 16S rRNA. Confocal laser scanning microscopy of larvae hybridized with the Cy5-labeled EUB388 III+, a specific probe for bacteria of the phylum Verrucomicrobia (*25*), showed that microcolonies (5-10 μm in diameter) consisting of several dozen cocci (500 nm in diameter) dispersed throughout the larvae’s interior (**Fig. 4A, B, Fig. S9A, D**). Staining with EUB388 I+, a universal probe for most bacteria excluding those in the phylum Verrucomicrobia (*25*), also revealed bacterial microcolonies of varying shapes and sizes (**Fig. S9C, F**).

**Fig. 4.**
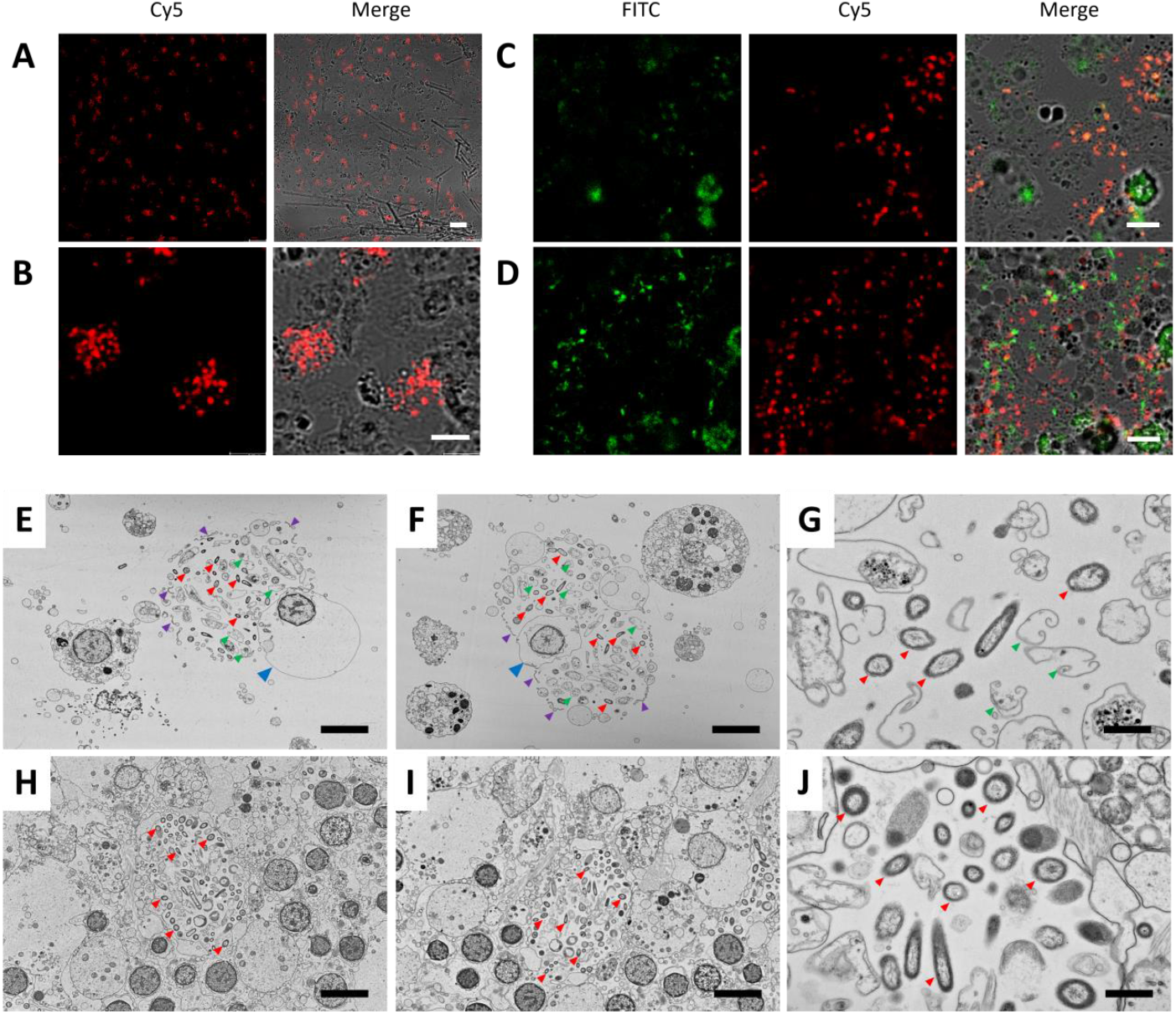
Visualization of symbiotic bacteria in early developmental stage of *M. nullarosette* by FISH and TEM. (A, B) Staining of larvae with Cy5-labeled EUB388 III+. Bacteria belonging to phylum Verrucomicrobia are specifically stained. (C) Double staining of larvae with FITC-labeled EUB388III+ and Cy5-labeled St-16S, a mixture of five specific probes designed for 16S rRNA of *Ca. S. tutelaris*. (D) Double staining of larvae with FITC-labeled EUB388 I+, a common probe for most eubacteria, and Cy5-labeled St-16S. (E-J) TEM images of the embryos (E-G) and larvae (H-J). Red arrows indicate live bacteria, green arrows indicate broken particles potentially lysed bacteria or organelles, blue arrows indicate host cells attached to the bacteriocyte-like structure, and purple arrows indicate disrupted membrane-like structures. Scale bars: 20 μm (A), 5 μm (B-D), 4.0 μm (E, F, H, I), and 0.8 μm (G, J).

To test whether the cocci stained with EUB388 III+ were indeed ‘Synechomycale', we performed double-staining experiments using EUB388 I+, EUB388III+, and ‘Synechomycale'-specific 16S rRNA probes (St-16S 1-5; **Table S9**). Initial staining with FITC-labeled EUB388 III+ and Cy5-labeled St-16S probes showed co-localized signals in the merged image, indicating that the cocci stained with EUB388 III+ were actually ‘Synechomycale’ (**Fig. 4C**). Double-staining with FITC-labeled EUB388 I+ and Cy-5-labeled St-16S probes showed that the fluorescent signals were present in the same area, but did not co-localize, indicating that ‘Synechomycale’ coexists with other bacteria in the microcolonies (**Fig. 4D**). Staining with FITC-labeled EUB388 I+ and Cy5-labeled EUB388 III+ showed similar results to the second experiment as expected (**Fig. S10A**). These results demonstrate that ‘Synechomycale’ is a spherical bacterium, approximately 500 nm in diameter, that forms microcolonies with other symbiotic bacterial species. These features closely resemble bacteriocytes, recognized in other sponges as habitats for symbiotic bacteria (*4, 24*). DAPI staining of sponge cell nuclei revealed their location near the bacteriocyte-like structures, although it was unclear whether they were inside or outside the host cells (**Fig. S11**).

### Mode of symbiosis of *Ca*. S. tutelaris in *M. nullarosette*

TEM images of embryos showed dozens of bacterial cells with varying sizes and shape clustered within small areas, approximately 10 μm in diameter (**Fig. 4E-G**). All bacteria-packed regions were surrounded by a membrane-like structure (purple arrows) and were invariably adjacent to the host cells, which were devoid of intracellular organelles except for nuclei. The membrane-like structure surrounding these clusters appeared disrupted, allowing bacteria to exist outside the host cell. Within the area, alongside intact bacterial cells, many particles lacking distinct morphology were observed, potentially representing lysed bacteria or other degraded materials. In larvae, host cells appeared more densely packed, while bacterial clusters were predominantly located in the extracellular matrix, surrounded by the host cells (**Fig. 4H-J**). These observations suggest that an extracellular symbiosis involving ‘Synechomycale’ and other bacteria commences from the embryonic stage.

## Discussion

The transmission modes of sponge symbionts, whether horizontal, vertical, or both, have long been debated, particularly since the discovery of diverse microorganisms within sponges (*4, 16*). Despite this interest, the origins and transmission modes of symbionts responsible for the production of bioactive metabolites remain largely unexplored. In this study, we advanced this field by revealing the relationship among mycalolides, their bacterial producer, and the host sponge *M. nullarosette* based on our establishment of its full lifecycle in aquaculture.

The complete genome sequence of the bacteria responsible for mycalolides production identified it as a novel genus within the phylum Verrucomicrobia, designated as *Candidatus* Synechomycale tutelaris. Surprisingly, the 16S rRNA of ‘Synechomycale’ shows only 89% identity to that of *Ca*. Epixenosomes, a symbiont of *Euplotidium arenarium*, the closest registered species, indicating the taxonomic uniqueness of the mycalolides producer. Despite the common presence of Verrucomicrobia bacteria in various environments (*26, 27*) and their frequent symbiotic relationship with aquatic organisms (*28*), the majority remain uncultured and are recognized as a typical group of the so-called ‘bacterial dark matter'. The genome of ‘Synechomycale’ harbors not only biosynthetic gene clusters for mycalolides but also for mycapolyols, with LC-MS confirming the presence of these metabolites in the sponge extract (**Fig. S6**). Notably, no other significant biosynthetic gene clusters were detected in the entire sponge metagenome, highlighting the bacterium’s prominent role in the host’s chemical defense. The presence of numerous mobile genetic elements suggested a significant degree of genome shrinking, with other symbionts and the host sponge presumably involved in the maintenance and proliferation of *?*Synechomycale*'*.Similarly, *Ca*. Endobryopsis kahalalidefaciens, a symbiotic bacterium producing kahalalides for the chemical defense of its host alga, lacks many genes associated with amino acid metabolism (*29, 30*), hindering its ability to exist independently.

Metagenomic and FISH analyses revealed that ‘Synechomycale’ are present throughout all developmental stages of *M. nullarosette*, forming bacteriocyte-like microcolonies alongside other symbionts. TEM images showed a potential transitional phase in embryos, where ‘Synechomycale’ moves from a temporary endosymbiosis state to an extracellular matrix, facilitating free proliferation (**Fig. 4**). We propose that phagocyte-like bacteriocytes capture a variety of symbionts before embryogenesis, incorporating them into early embryos to establish vertical transmission. This mechanism ensures the secure and efficient transmission of symbiotic consortia, critical for subsequent generations (**Fig. 5**).

**Fig. 5.**
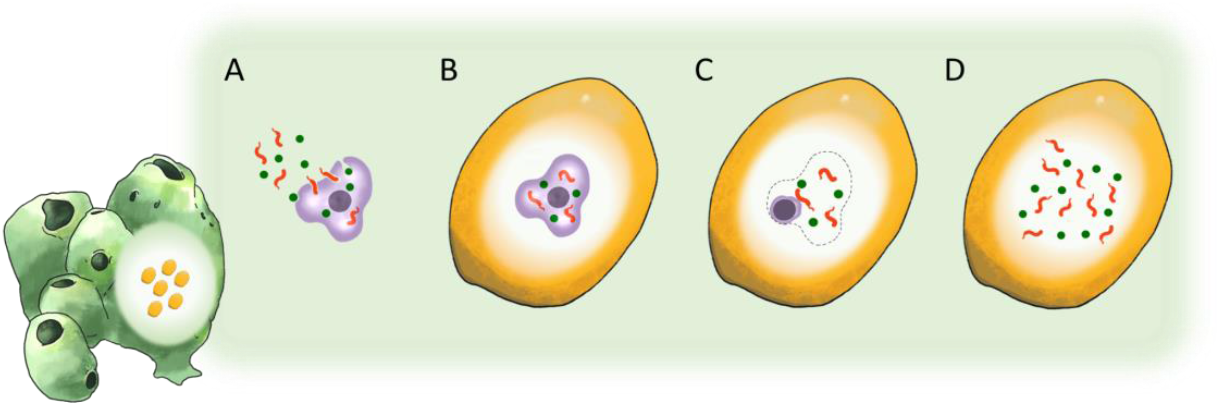
Hypothetical image of the *Ca*. S. tutelaris vertical transmission through bacteriocytes. (A) Symbiotic bacteria are ingested by bacteriocyte in the adult sponge. (B) The cells move into the early embryo. (C) The cell membrane is disrupted. (D) Symbiotic bacteria spread out in the larvae. Observations of stages C and D allow us to infer the earlier processes described in A and B.

Recent reviews have summarized various vertical transmission strategies in sponges, including bacteriocyte- or nurse cell-mediated transfer, phagocytosis by oocytes, and bacterial self-migration into embryos (*16, 24*). The bacteriocyte-like structures observed in ‘Synechomycale’ closely resembles those in *Svenzea zeai* (*31*) and *Petrosia ficiformis* (*32*), where bacteria are housed in vacuole-like cavities lacking intracellular organelles (*32, 33*). While the prior study reported incomplete vertical transmission, with larvae inheriting only 44.8% of parental microbes (*34*), our data demonstrates a more faithful transmission of symbionts in *M. nullarosette*. Bacteriocyte- or nurse cell-mediated vertical transmission appears more effective for transferring a ‘bacterial package’ compared to self-migration or phagocytosis by oocytes.

The sponge *M. nullarosette* appears to inherit its entire symbiont consortium, including cytotoxin-producing and auxiliary bacteria, through bacteriocyte-mediated vertical transmission. This efficient and reliable delivery system is especially critical for symbionts lacking essential genes to survive independently outside the host environment. The *Mycale*-Synechomycale symbiosis provides a compelling model for primitive mutualism between multicellular hosts and unicellular symbionts. Thus, certain sponges have evolved an efficient mechanism to guarantee the transmission of vital bacteria involved in their chemical defense, which has contributed to their survival throughout their long history.

## Supporting information

Supplementary_materials_mycalolideVT

## Acknowledgments

We thank Drs. Gota Goshima and Hitoshi Sawada of Nagoya University for invaluable help with sample collection. We also thank Dr. Kensuke Igarashi of the National Institute of Advanced Industrial Science and Technology for discussion on TEM observation.

## Authors contributions

Conceptualization: M.F., K.T.; Methodology: M.F., Y.I., M.F., N.K., K.Y.; Investigation: M.F., Y.I., M.F., A.N., K.Y., K.T.; Funding acquisition: M.F., S.M., R.S, K.T.; Project administration: M.F., K.T.; Writing – original draft: M.F., K.T.; Writing – review & editing: M.F., Y.I., M.F., N.K., A.N., K.Y., S.M., R.S., K.T

## Data availability

Nucleotide sequence data reported are available in the DDBJ Sequenced Read Archive under DDBJ with the accession number DRA010842 and DRA011078 for metagenomic data, and AP024111 and AP024164 for the complete circular genomes of *Ca*. S. tutelaris. Data is provided within the manuscript or a supplementary information file.

