## Supplementary_materials_mycalolideVT for "Symbiotic Bacterial Partners for Chemical Defense are Vertically Transmitted in a Marine Sponge"

**The PDF file includes:**

Materials and Methods
Figs. S1 to S11
Tables S1 to S9
References

### Materials and Methods

Collection of the marine sponge *Mycale nullarosette*

Specimens of *M. nullarosette* were collected by hand along the shores of Amakusa Islands in Kumamoto Prefecture (32º 31′N, 130º 25′E), and Suga Island in Mie Prefecture (34º 48′N, 136º 87′E) during the low-tide periods annually from May to June annually from 2016 to 2020.

**Embryos:** Specimens were dissected into 5 mm pieces and then immersed in $Ca^{2+}$- and $Mg^{2+}$-free artificial seawater (CMF, 449 mM NaCl, 9 mM KCl, 33 mM $Na_2SO_4$, 2.15 mM $NaHCO_3$, 10 mM Tris-HCl pH 8.0, 2.5 mM ethylene glycol tetraacetic acid pH 8.0) for 5 minutes. Subsequently, the sponge specimens were transferred to fresh CMF and gently shaken to lyse the extracellular matrix. This process was repeated until the embryos dissociated from the sponge specimens. The detached embryos were collected in microcentrifuge tubes and thoroughly rinsed several times with filter-sterilized seawater.

**Larvae:** Mature *M. nullarosette* individuals were transferred into the mesh bags (0.2 mm mesh) and kept in the sea overnight. Hatched swimming larvae, approximately 0.5 mm in diameter, were collected from the mesh bags and concentrated via filtration (pore size: 40 μm). The collected larvae were then transferred into microcentrifuge tubes and washed several times with filter-sterilized seawater.

Aquaculture of *M. nullarosette* throughout the life cycle

Collected larvae were placed in plastic Petri dishes and left for one week to settle on the bottom of the dishes. The dishes were then divided into three groups and placed in distinct environments: an indoor tank, an outdoor tank at a depth of 1 meter, and a shaded cage in the sea at a depth of 2 meters, respectively. In the indoor tank, seawater was replaced with natural seawater three times a week, and the larvae were fed daily with phytoplankton Phytogold-S (Brightwell Aquatics, Alabama, USA). In the outdoor tank, natural seawater pumped from the sea was continuously added. Juvenile sponges grown in these three environments, approximately six months old, were collected.

Preparation of metagenomic DNA

Specimens were gently squeezed by hand immediately after collection and stored at 4°C in sterilized bottles for transportation to the laboratory (10 min). The cell suspension (1.5 mL × 7 tubes) was immediately fractionated by stepwise centrifugation at 1, 2, 3, 4, 5, 6, and 12 × $10^3$ rpm for 1 min at each step, resulting in seven precipitated fractions. Genomic DNA was extracted from each precipitate using the QIAamp DNA Stool Mini Kit (Qiagen, Hilden, Germany), following the manufacturer's protocol. Metagenomic DNA from embryos or larvae was also extracted using the same procedure.

Metagenome analyses for sponge-derived metagenomic DNA

Metagenomic DNA was analyzed by Illumina HiSeq 100 bp pair-end sequencing, MiSeq 300 bp pair-end sequencing, and MinION long read sequencing using Ligation Sequencing Kit 1D (Oxford Nanopore Technologies, Oxford, UK). Additional NGS sequencing was performed on metagenomics DNA prepared from sponge samples collected in different years and from various sampling sites, using the same protocols. The sequencing data was deposited in the DDBJ with the accession number DRA010842 for the Amakusa samples and DRA011078 for the Suga Island samples.

HiSeq and MiSeq read datasets were assembled using the Velvet assembler (v1.2.10, hash length 35 bases) (*35*), resulting in approximately 5,000 contigs (each > 500 bases). To close gaps between contigs, DNA fragments were PCR amplified using metagenomic DNA as a template (Table. S2). The amplicons were digested by restriction enzymes and ligated into the pBCSK+ cloning vector (Agilent, CA, USA), then transformed into NEB 10-beta *E. coli* competent cells (New England Biolabs, MA, USA) via electroporation. Plasmids extracted from the transformants with the AxyPrep Plasmid Miniprep Kit (Axygen Biosciences, CA, USA) were subjected to the capillary sequencing using the BigDye Terminator v3.1 Cycle Sequencing Kit (Thermo Fisher, MA, USA), following the manufacturer's protocols. Sequence information for binning analysis was obtained using Jemboss. (*36*) Long-read datasets derived from MinION sequencing were assembled by Flye (v2.7) (*37*) to generate draft circular genome sequence. The circular genomes were further refined to correct miss-called bases using short-read data with bowtie2 (v2.3.4.2) (*38*) and pilon (v1.23) (*39*), resulting in the complete circular genomes of the mycalolide producer, *Ca*. S. tutelaris. The accession number are AP024111 for the Amakusa strain and AP024164 for the Suga Island strain. Detection of the secondary metabolites biosynthetic gene clusters was performed using antiSMASH bacterial version (v.6.0).(*40*) Detection of the coding sequences (CDSs) and functional annotation were performed using RAST (v.2.0),(*23*) and DFAST (v. 1.3.0).(*41*)

Taxonomic analyses of *Ca*. S. tutelaris in the phylogenetic tree of 16S rRNA

To construct taxonomic trees of the 16S rRNA of bacteria within the phylum Verrucomicrobia, we selected 29 bacterial strains comprising eight genera within the class Opitutae, and 25 bacterial strains belonging to other classes within the Phylum. The 16S rRNA gene sequences of these selected bacteria were retrieved from the SILVA rRNA database. (*42*) These sequences were aligned using MAFFT (Version 7.490) with the 'Auto' algorithm option and a scoring matrix set at 200PAM/k=2. The aligned sequences were then used to construct a phylogenetic tree using MrBayes (Version 3.2.6). The analysis utilized the GTR substitution model with the following MCMC parameters: a chain length of 100,000, four heated chains, a heated chain temperature of 0.2, a subsampling frequency of 200, and a burn-in length of 90,000.

LC-MS of the mycalolides in each life stage

Mature sponge specimens, embryos, larvae, and juvenile sponges were extracted with MeOH overnight at room temperature. The extracts were analyzed by LC-MS with $H_2O$-MeCN gradient system using reversed-phase $C_{18}$ column (Cosmosil MSII 2.5$C_{18}$ 2.0 mm i.d. × 100 mm). Ions corresponding to three major mycalolides, mycalolide A (**1**: *m/z* 909), mycalolide B (**2**: *m/z* 1027), and mycalolides C (**3**: *m/z* 997) were extracted.

Bacterial consortia analysis in each life stage

The sequence of V3-V4 regions in 16S rRNA was amplified from metagenomic DNA using conditions recommended by Illumina. The amplicons were then purified by AxyPrep PCR Clean-Up Kit (Axygen Biosciences, CA, USA) or MagExtractor™ (TOYOBO, Osaka, Japan). The purified amplicon was analyzed by MiSeq 300 bp pair-end sequencing. Taxonomy analysis was performed using Qiime2 (ver. 2021.4) with SILVA database (ver. 138.1) (*42*) to afford taxonomic annotation to species level.

Detection of the *myl* gene cluster in each life stage and environmental DNA

To detect the *myl* gene cluster, specific primers were designed for three parts of the *myl* cluster (Table S8). These sequences were amplified from environmental DNA including seawater-derived metagenomes, bottom sediment-derived metagenomes, and neighboring sponge-derived metagenomes. The preparation of each metagenome is described below. The amplicons were analyzed by agarose gel electrophoresis and visualized with ethidium bromide.

Seawater-derived metagenomes: The seawater (1 L) collected from the habitat of *M. nullarosette* was serially filtered by 10 μm, 0.45 μm, and 0.1 μm pore size membranes. The 0.45 μm and 0.1 μm membranes were placed in the 15 mL tubes and vigorously shaken with 1.0 mL of filter-sterilized seawater to wash out the filtrates, then centrifuged to obtain cell pellets. The pellets were combined and subjected to genomic DNA extraction in the same manner as described above.

Bottom sediment-derived metagenomes: Bottom sediment (20 mL in 50 mL tubes) collected from the habitat of *M. nullarosette* was vigorously shaken with 20 mL of filter- sterilized seawater containing 0.2% Triton-X100 and left to stand for 3 min. The upper suspension was decanted into new tubes and centrifuged to obtain pellets. The pellets were combined and subjected to genomic DNA extraction in the same manner described above.

Neighboring sponge-derived metagenomes: A small piece of unidentified sponge specimen (most likely *Halichondria okadai*), located within 1 m of *M. nullarosette*, was collected by hand and immediately squeezed to obtain symbionts suspension. This was centrifuged, and the resulting cell pellet was subjected to genomic DNA extraction in the same manner as described above.

Fluorescent *in situ* hybridization for embryos and larvae

Embryos and larvae (50 embryos or 30 larvae in 50 mL of seawater) were jellified by addition of iPGell (Genostaff, Tokyo, Japan) according to the manufacturer's protocol. The jellified samples were fixed with G-fix fixation solution (Genostaff) at 4 $^{o}$C. Paraffin sections (5 mm thickness) of embryos and larvae for *in situ* hybridization were prepared.

Paraffin-embedded sections of *M. nullarosette* embryos and larvae were deparaffinized by soaking in xylene for 5 min twice, followed by a series of EtOH rinses at concentrations of 100%, 95%, 90%, 80%, 70%, and 50% EtOH for 5 min each. The sections were then washed with distilled water and phosphate-buffered saline (PBS) for 5 min each. Deparaffinized slides were treated with proteinase K (10 μg/mL in PBS containing 0.1% Tween 20) at 37 $^{o}$C for 15 min, then rinsed with a glycine solution (2 mg/mL in PBS containing 0.1% Tween 20) at room temperature for 5 min. The slides were washed twice with PBS for 5 min each, re-fixed with 4% paraformaldehyde in PBS for 10 min, then washed again twice with PBS for 5 min. Sections were covered with hybridization buffer [20% formamide, 4× SSC (150 mM NaCl, 15 mM trisodium citrate, pH 7.0), 0.02% SDS, 1 mg/mL yeast tRNA, 1× Denhardt's solution, 5 mM EDTA, 10 % dextran sulfate] and incubated at 42 $^{o}$C for 30 min. Fluorescence-labeled (Cy5 or FITC) oligo DNA probes in hybridization buffer (total 10 pmol/100 μL) were applied to the sections and incubated at 42 $^{o}$C for 3 hours in a humidified chamber (probe sequences: see Table S9). Slides were washed with 2× SSC for 5 min at room temperature, followed by 0.5× SSC at 42 $^{o}$C for 10 min. They were then washed with Tris-buffered saline [TBS: 50 mM Tris-HCl (pH

7.5), 150 mM NaCl] for 5 min twice. Finally, the slides were air-dried at room temperature, mounted with ProLong Diamond Antifade Mountant (Thermo Fisher), and observed with a confocal laser scanning microscope (Leica TCS-SP5, Wetzlar, Germany).

Transmission electron microscopy (TEM) analyses of embryos and larvae

Embryos and larvae were fixed in 2% glutaraldehyde in PBS (Electron Microscopy Science, PA, USA). The samples were subsequently post-fixed in 2% osmium tetroxide (Heraeus Chemicals South Africa, South Africa) for 2 hours in an ice bath. The specimens were dehydrated in a graded series of ethanol concentrations (30, 50, 70, 90, 100, 100, 100 % for 15 min each), and embedded in epoxy resin (TAAB Laboratories, UK). Ultrathin sections were obtained using an ultramicrotom. The sections, stained with uranyl acetate for 15 minutes and a lead staining solution for 5 minutes, were examined with a TEM at 100 kV (Hitachi H-7600, Tokyo, Japan).

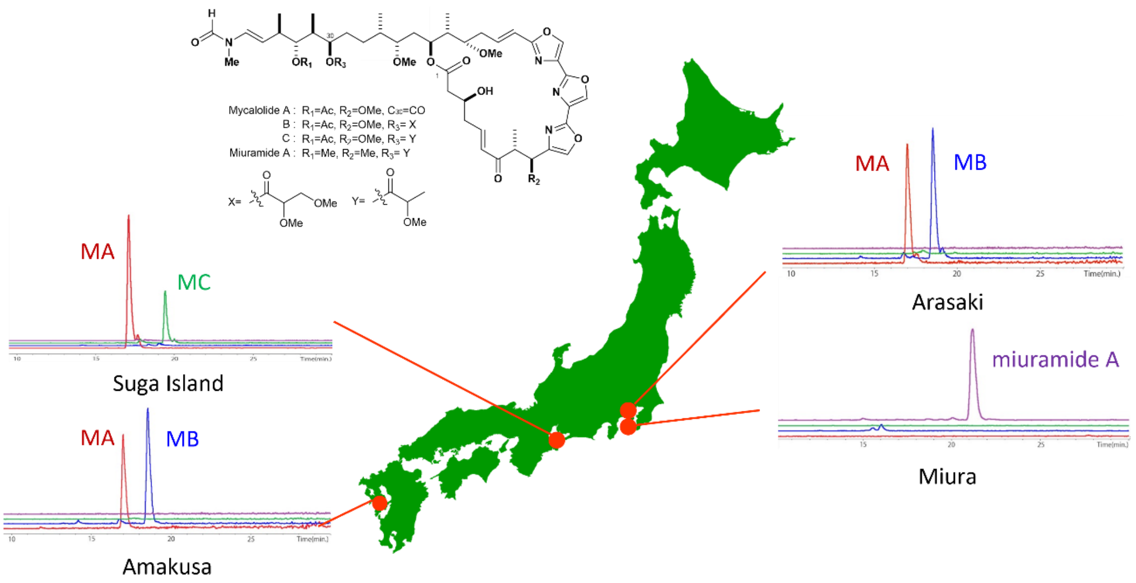


**Fig. S1. Chemical profiles of *Mycale* sponges from four collection sites in Japan: Amakusa, Suga Island, Arasaki, and Miura.** The distance between Arasaki and Miura is only approximately 5 km. MA, MB, and MC represent mycalolides A, B, and C, respectively.

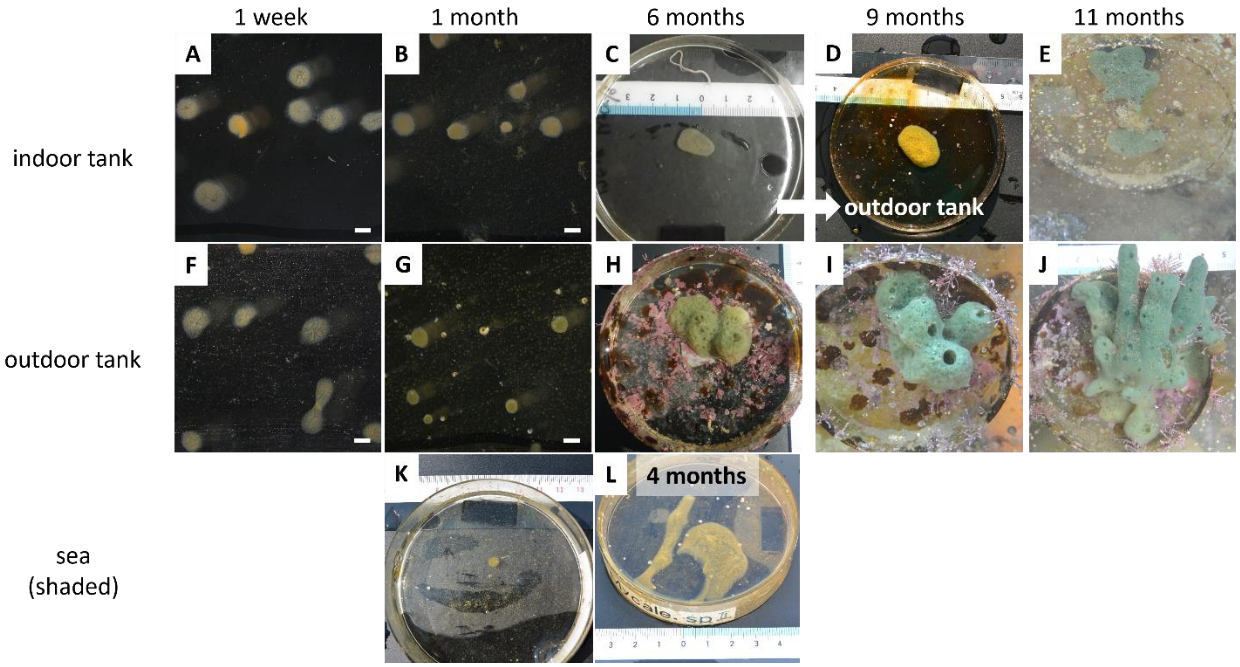


**Fig S2. Cultivation of the marine sponge *Mycale nullarosette*.** (A-C) Juvenile sponges cultured in an indoor tank after settlement: (A) one week, (B) one month, and (C) six months. (D-E) The Petri dish was transferred to an outdoor tank: (D) nine months and (E) eleven months. (F-J) Juvenile sponges cultured in the indoor tank after the settlement: (F) one week, (G) one month, (H) six months,(I) nine months, and (J) eleven months. (K-L) Juvenile sponges cultured in a shaded cage placed in the sea: (K) one month and (L) four months. Scale bar represents 1 mm.

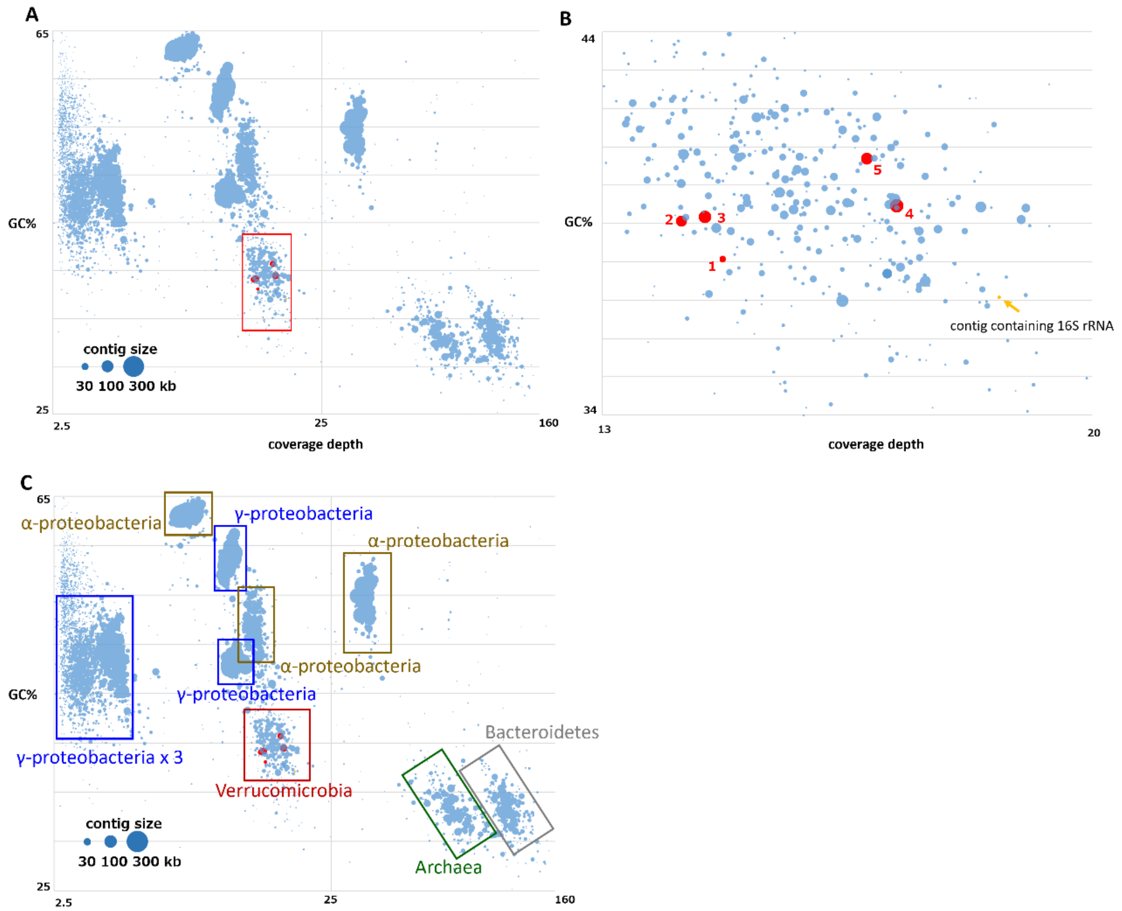


**Fig. S3. Binning analysis of assembled contigs based on their coverage depth and GC content.** (A) Distribution of all contigs derived from metagenomic analyses by Illumina Hiseq, (B) Close-up view of the bin containing the mycalolides biosynthetic genes. (C) Phylum-level classification of each bin based on the genes they contain. The size of each circle indicates the contig size. Red circles represent contigs containing *myl* biosynthetic genes. A yellow circle represents the contig containing 16S rRNA.

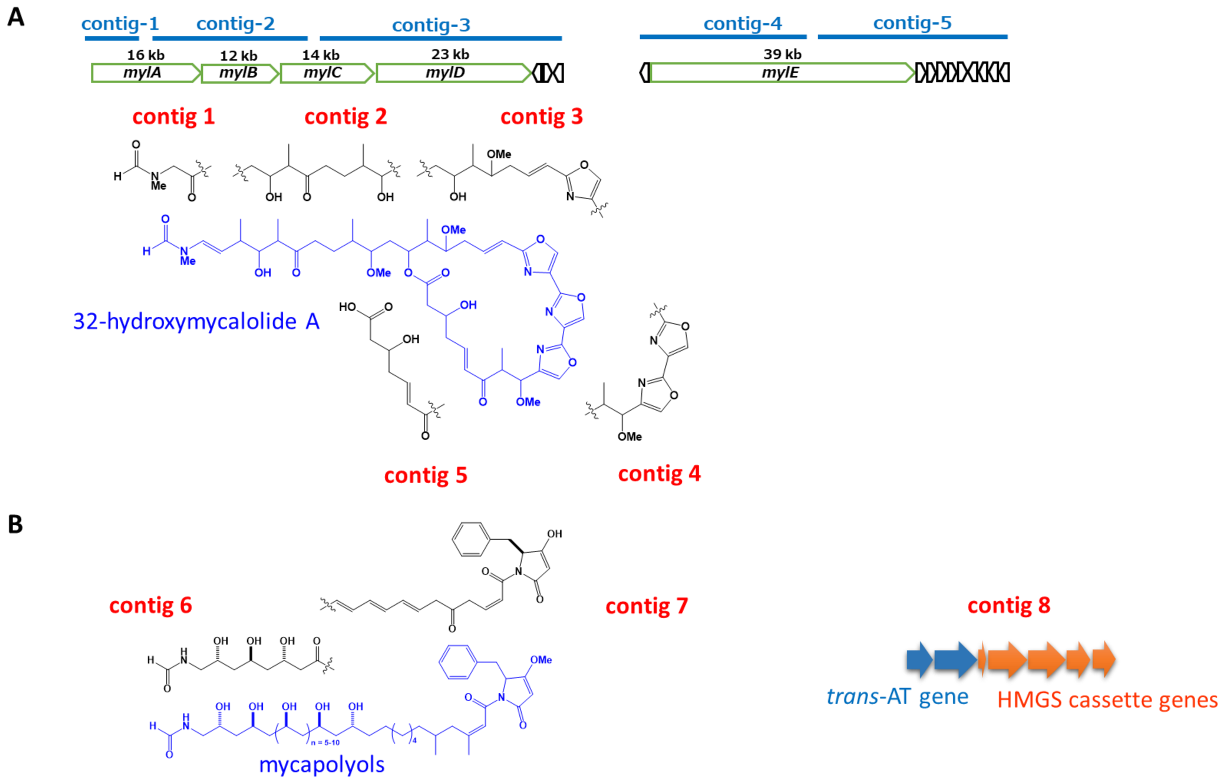


**Fig. S4**. **Summary of the predicted structures of contigs containing PKS-NRPS genes.** (A) Contigs 1-5 are involved in mycalolides biosynthesis. (B) Contigs 6-8 are involved in mycapolyols biosynthesis.

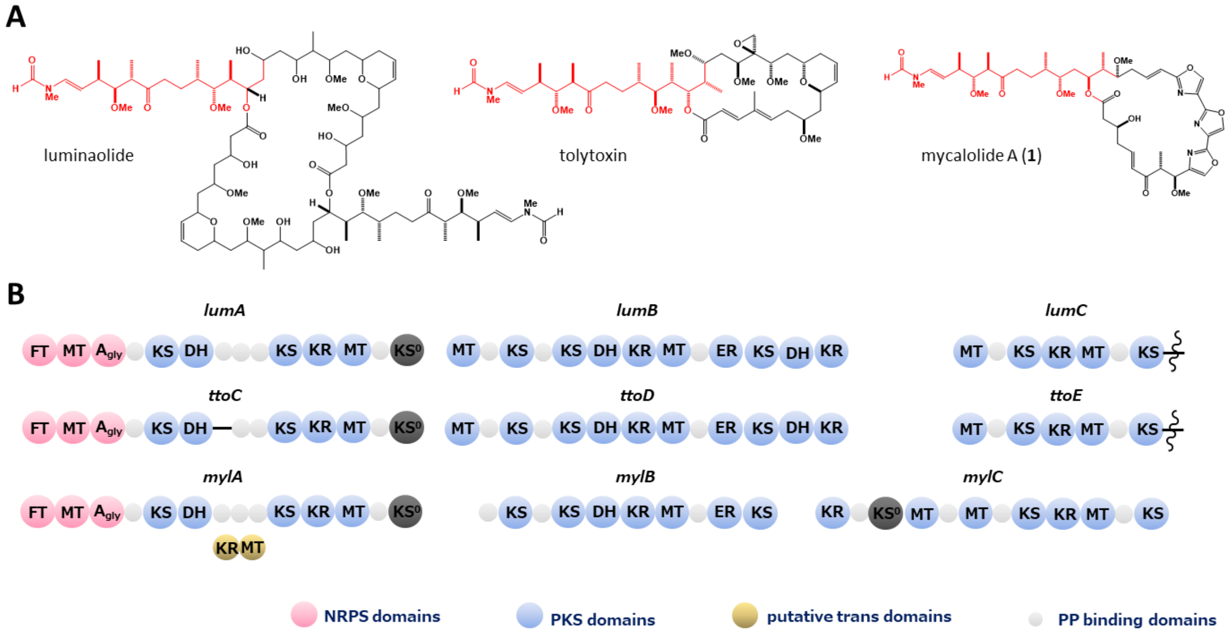


**Fig. S5. Comparison of biosynthetic gene clusters for common substructures in three actin depolymerizing macrolides.** (A) Structures of three actin depolymerizing macrolides, with the common side chain highlighted in red. (B) Domain architecture of the biosynthetic gene clusters responsible for the side chain parts in cyanobacteria-derived luminolide (*lum*), tolytoxin (*tto*), and sponge-derived mycalolides (*myl*).

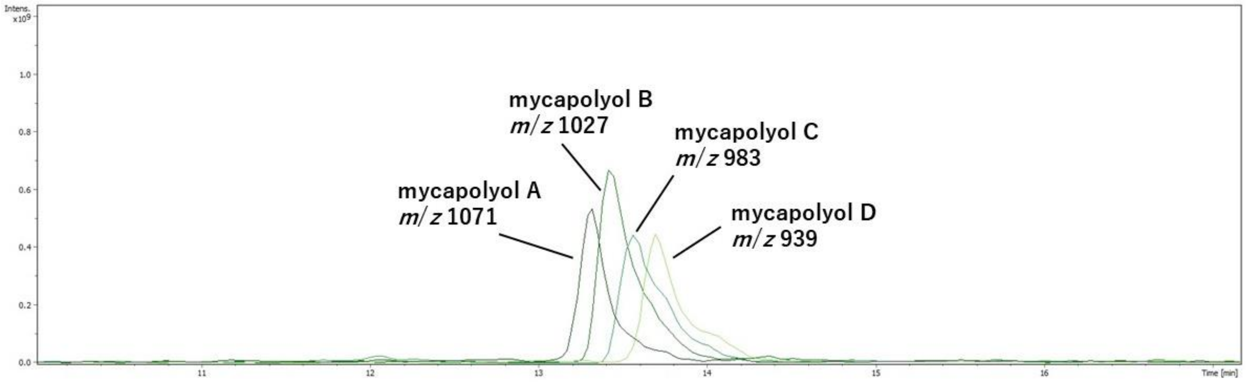


**Fig. S6. LC-MS data of the organic extract from the adult sponges.** Four ion peaks are displayed: *m*/*z* 1071 for mycapolyol A, *m*/*z* 1027 for mycapolyol B, *m*/*z* 983 for mycapolyol C, *m*/*z* 939 for mycapolyol D.

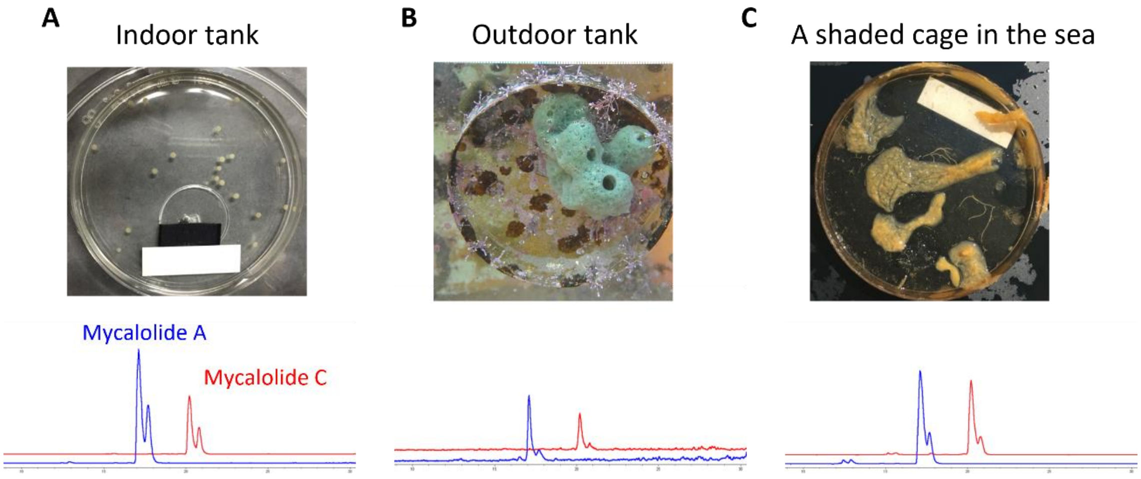


**Fig. S7. LC-MS data of the organic extract from juvenile sponges.** LC-MS data for the sponges grown in different environments: (A) indoor tank, (B) outdoor tank, and (C) shaded cage in the sea.

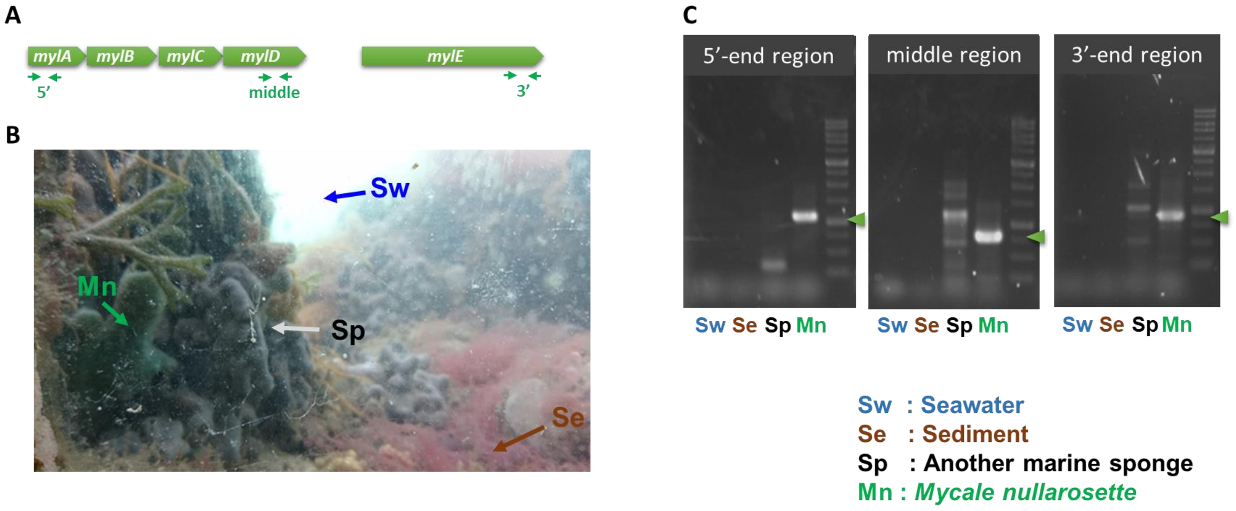


**Fig. S8. PCR amplification using specific primers for the *myl* gene cluster. (**A) Target regions for specific primers to be amplified. The primers were designed in the 5′-end, middle, and 3′-end regions of the *myl* gene cluster. (B) Photograph of the sample collection area. Metagenomic DNA was prepared from surrounding seawater (Sw), sediment (Se), a different marine sponge (Sp) and *Mycale nullarosette* (Mn). (C) Results of agarose gel electrophoresis for PCR amplification using metagenomic DNA as templates. Green arrows indicate the expected product size for designed primers.

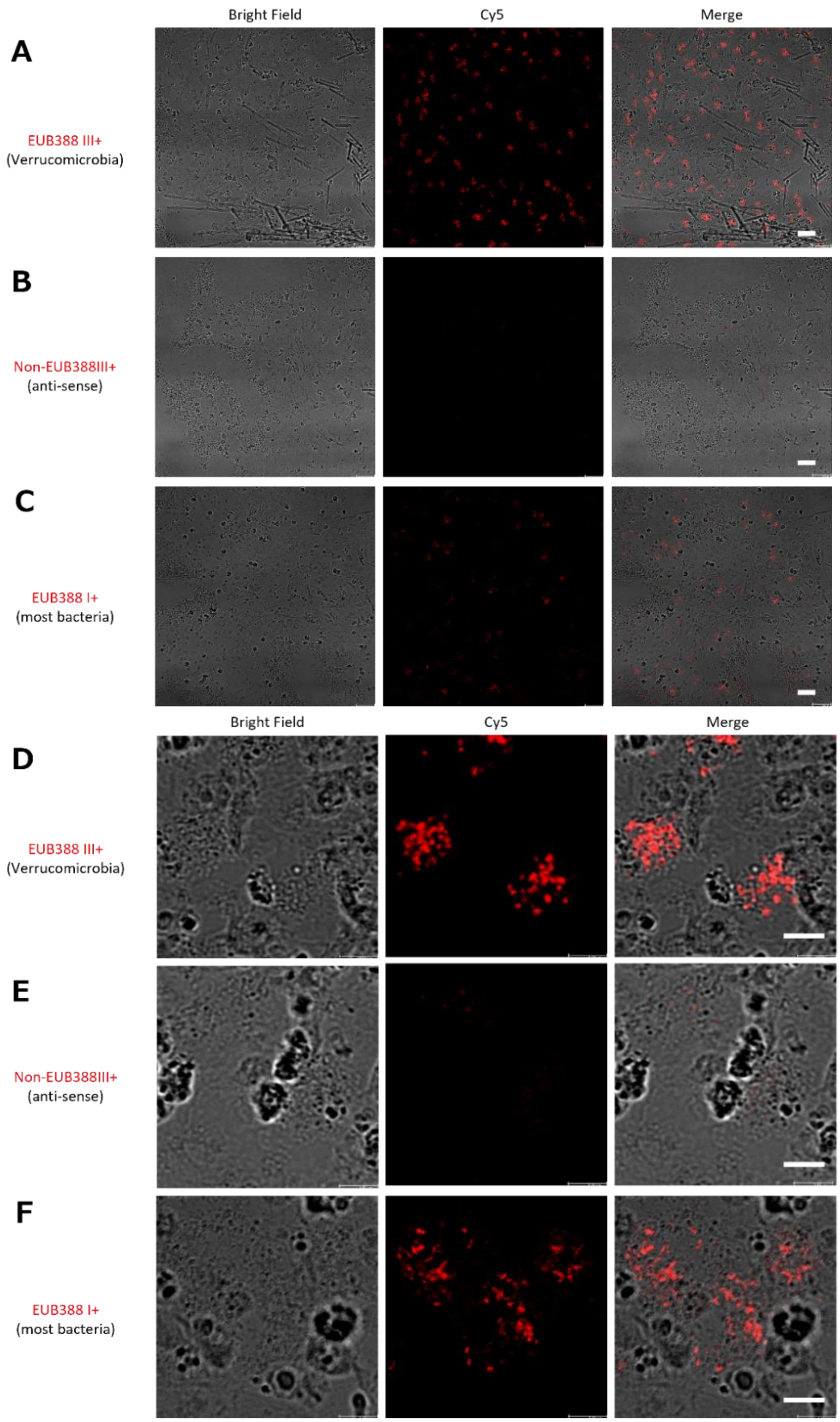


**Fig. S9. Fluorescence in situ hybridization images of larvae of *M. nullarosette.*** (A, D) EUB388 III+, a specific probe for bacteria belonging to Phylum Verrucomicrobia. (B, E) Non-EUB388 III+, an anti-sense probe of EUB388 III+. (C, F) EUB388 I+, a universal probe for most eubacteria. Scale bars represent 20 μm for A-C, and 5 μm for D-F.

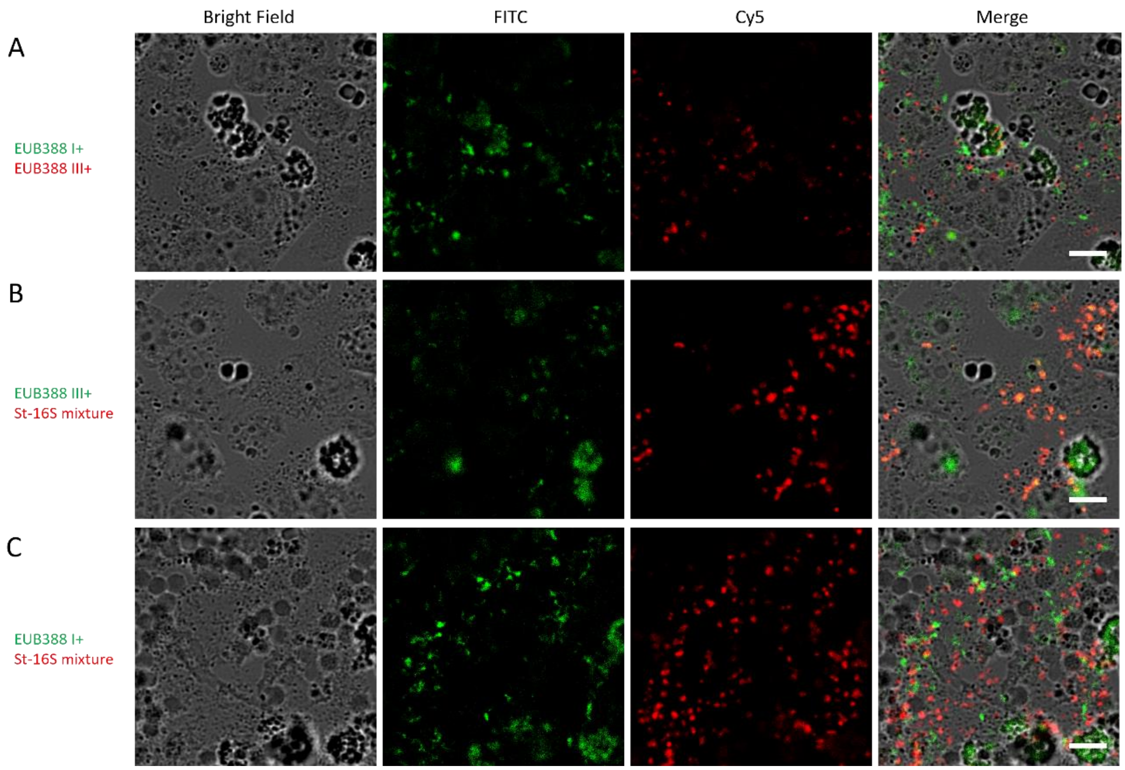


**Fig. S10. Fluorescence in situ hybridization images of larvae of *M. nullarosette.*** A double staining for (A) EUBI+ and EUB388III+, (B) EUB388III+ and St-16S, a mixture of five specific probes designed based on 16S rRNA of *Ca. S. tutelaris*. (C) EUB388 I+ and St-16S. Scale bars represent 5 μm.

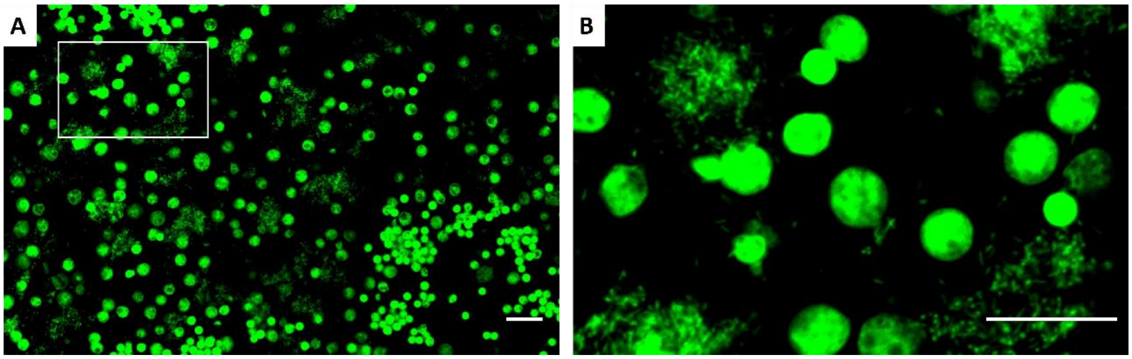


**Fig. S11. DAPI staining of larvae observed by confocal laser scanning microscopy.** (A) Whole image: A comprehensive view of the larvae, highlighting the overall staining pattern of sponge cell-derived nuclei and bacteria-derived nuclei within the larval. (B) Enlarged image: A magnified view of a specific area indicated in panel A. Bacteriocyte-like areas are adjacent to the host cell nuclei. Scale bars represent 10 µm.

**Table S1. Contigs containing PKS-NRPS genes.**

| | Length (bp) | Description |
|---|---|---|
| contig 1 | 8201 | PKS-NRPS hybrid (Gly) |
| contig 2 | 23590 | PKS |
| contig 3 | 30953 | PKS-NRPS hybrid (Ser) |
| contig 4 | 24325 | PKS-NRPS hybrid (Ser x 2) |
| contig 5 | 26558 | PKS |
| contig 6 | 22805 | PKS-NRPS hybrid (Gly) |
| contig 7 | 28346 | PKS-NRPS hybrid (Phe) |
| contig 8 | 18458 | transAT and HMGS cassette |
| contig 9 | 12371 | PKS |
| contig 10 | 19361 | PKS |

**Table S2. PCR Primers used for closing the contigs gap.**

| Position | Primer Name | Sequence |
|---|---|---|
| Contigs 1 to 2 | Gap-F1 | TTCTGCACCCCTGCCTTATG |
| | Gap-R2 | AGTACCGAGGGACAGACTCC |
| Contigs 2 to 3 | Gap-F2 | CTTGGAAGCTGCATCTGGGA |
| | Gap-R3 | TCCAACGTGTTTCAGGCACT |
| Contigs 4 to 5 | Gap-F4 | AAGAAGGTGCTGGGTCGATG |
| | Gap-R5 | GGGCTTTAAGTCGCTCCCAT |

**Table S3. Common motif sequences in KS domains.**

| | Common Motif 1 | Common Motif 2 | Common Motif 3 | Predicted Function |
|---|---|---|---|---|
| KS1 | CSSSL | HGTGT | NIGH | active |
| KS2 | CSSSL | HGTGT | HTGH | active |
| KS3 | CSSSL | SGVVD | HTGH | inactive |
| KS4 | CSSSL | HGTGT | NLGH | active |
| KS5 | CSSSL | HGTGT | NIGH | active |
| KS6 | CSSSL | HGTGT | NIGH | active |
| KS7 | CASSL | QAVGS | SIGH | inactive |
| KS8 | CSSSL | HGTGT | NIGH | active |
| KS9 | CSSSL | HGTGT | NIGH | active |
| KS10 | CSSSL | QALGS | NIGH | inactive |
| KS11 | CSSAL | HGTGT | NVGH | active |
| KS12 | CSSSL | AANGS | NLGH | inactive |
| KS13 | CSSTL | AATGS | NIGH | inactive |
| KS14 | CSSSL | ATNGS | NIGH | inactive |
| KS15 | CASAL | ATNGS | NIGH | inactive |
| KS16 | CSSTL | AANGS | NIGH | inactive |
| KS17 | CSSSL | HGTGT | NIGH | active |
| KS18 | CSSSL | QTLGS | NIGH | inactive |
| KS19 | CSSSL | HGTGT | NIGH | active |
| KS20 | CSSSL | HGTGT | NMGH | active |
| KS21 | CSSSL | HGTGT | NVGH | active |
| KS22 | CPSSY | HGTAS | NIGH | active |

Sequences shown in red letters were not conserved suggesting that these domains are malfunctional $KS^0$.

**Table S4. General genomic information and results of in silico DNA-DNA hybridization.**

| Family | Genus | Species | Strain | Genome Size | GC% | Accession | ANI | Tetra | DDH |
|---|---|---|---|---|---|---|---|---|---|
| Puniceicoccaceae | *Ca*. Synecomycale | tutelaris | Am1 | 2.53 Mb | 40.7 | AP024111 | na | na | na |
| Puniceicoccaceae | *Ca*. Synecomycale | tutelaris | Sg1 | 2.52 Mb | 40.7 | AP024164 | 99.8 | 0.999 | 98.5 |
| Puniceicoccaceae | *Coraliomargarita* | *akajimensis* | DSM45221 | 3.75 Mb | 53.6 | NC014008.1 | 64.4 | 0.492 | 26.2 |
| Akkermansiaceae | *Akkermansia* | *muciniphila* | JCM30893 | 2.89 Mb | 55.6 | AP021898.1 | 64.0 | 0.452 | 20.6 |
| Methylacidiphilaceae | *Methylacidiphilum* | *infernorum* | V4 | 2.29 Mb | 45.5 | NC010794.1 | 63.9 | 0.655 | 21.6 |

### Table S5. Annotation results of bacterial genomes belonging to the phylum Verrucomicrobia

| Species | | *Ca.* S. tutelaris | *C. akajimensis* | *A. muciniphila* | *M. infernorum* |
|---|---|---|---|---|---|
| Strain | | Am1 | DSM 45221 | JCM 30893 | V4 |
| Genome Size | | 2.53 Mb | 3.75 Mb | 2.89 Mb | 2.29 Mb |
| Total CDS | | 2987 | 3081 | 2323 | 2572 |
| Subsystems | | 162 | 251 | 272 | 267 |
| RNAs | | 51 | 50 | 59 | 49 |
| CDS in subsystem | Cofactors, Vitamins, Prosthetic Groups, Pigments | 60 | 101 | 170 | 146 |
| | Cell Wall and Capsule | 4 | 32 | 81 | 40 |
| | Virulence, Disease and Defense | 20 | 15 | 31 | 44 |
| | Potassium metabolism | 1 | 13 | 3 | 10 |
| | Photosynthesis | 0 | 0 | 0 | 0 |
| | Miscellaneous | 8 | 19 | 21 | 4 |
| | Phages, Prophages, Transposable elements, Plasmids | 2 | 1 | 2 | 0 |
| | Membrane Transport | 20 | 30 | 23 | 21 |
| | Iron acquisition and metabolism | 0 | 1 | 17 | 0 |
| | RNA Metabolism | 34 | 105 | 122 | 106 |
| | Nucleosides and Nucleotides | 37 | 54 | 50 | 66 |
| | Protein Metabolism | 59 | 153 | 184 | 147 |
| | Cell Division and Cell Cycle | 0 | 24 | 11 | 29 |
| | Motility and Chemotaxis | 0 | 3 | 0 | 0 |
| | Regulation and Cell signaling | 2 | 10 | 9 | 6 |
| | Secondary Metabolism | 7 | 4 | 4 | 5 |
| | DNA Metabolism | 33 | 50 | 92 | 39 |
| | Fatty Acids, Lipids, and Isoprenoids | 18 | 49 | 61 | 55 |
| | Nitrogen Metabolism | 7 | 23 | 3 | 13 |
| | Dormancy and Sporulation | 1 | 3 | 1 | 1 |
| | Respiration | 45 | 75 | 53 | 84 |
| | Stress Response | 10 | 41 | 32 | 32 |
| | Metabolism of Aromatic Compounds | 1 | 4 | 1 | 1 |
| | Amino Acids and Derivatives | 133 | 196 | 205 | 185 |
| | Sulfur Metabolism | 11 | 52 | 24 | 10 |
| | Phosphorus Metabolism | 6 | 31 | 32 | 31 |
| | Carbohydrates | 44 | 169 | 129 | 162 |

| | | | | | |
|---|---|---|---|---|---|
| not in subsystem | Transposase | 102 | 0 | 0 | 0 |
| | Mobile Elemental Protein | 124 | 0 | 0 | 0 |
| | Transposase OrfB | 39 | 0 | 0 | 2 |
| | IS1478 Transposase | 6 | 0 | 0 | 0 |
| | Integrase Catalytic Domain | 20 | 0 | 0 | 0 |
| | Transposase IS111A/IS1328/IS1533 | 0 | 4 | 0 | 0 |
| | Transposase, IS605 OrfB | 0 | 0 | 0 | 2 |
| | Transposon IS605 OrfA, integrase-resolvase | 0 | 0 | 0 | 5 |
| | Transposon IS605 OrfB | 0 | 0 | 0 | 6 |
| | Transposon related ORF | 0 | 0 | 0 | 1 |
| | Integrase | 0 | 0 | 3 | 0 |
| | Integration host factor beta subunit | 0 | 0 | 2 | 0 |
| in subsystem | hypothetical | 9 | 20 | 34 | 21 |
| not in subsystem | hypothetical | 1820 | 1405 | 861 | 1167 |

**Table S6. PKS-NRPS related biosynthetic gene clusters in *Ca*. S. tutelaris genomes.**

| | type | description | Am1 | Sg1 |
|---|---|---|---|---|
| Cluster 1 | transAT-PKS-NRPS-like | *mylA-D* | 65882 | 65885 |
| Cluster 2 | transAT-PKS-NRPS-like | *mylE* | 39426 | 39426 |
| Cluster 3 | transAT-PKS-NRPS-like | *mypA* | 18342 | 18342 |
| Cluster 4 | type 1 PKS-NRPS-like | *mypB-C* | 27549 | 27549 |
| Cluster 5 | PKS-like | *mypD-J* | 6728 | 6729 |
| Cluster 6 | transAT-PKS-like | Function unknown | 13470 | nd |
| Cluster 7 | type 3 PKS | HMG-CoA synthase | 1176 | 1176 |
| Cluster 8 | transAT-PKS-like | potentially inactive | 48401 | 48424 |

nd: not detected.

**Table S7. Microbial consortia in the second generation of *Mycale nullarosette* grown in an outdoor tank and a shaded cage in the sea.**

| Phylum | Class | Genus | Species | ratio % |
|---|---|---|---|---|
| Second generation sponge_Outdoor tank | | | | |
| Proteobacteria | Alphaproteobacteria | *Hyphomonas* | | 29.0 |
| Proteobacteria | Alphaproteobacteria | *Hyphomicrobium* | *aestuarii* | 26.9 |
| Crenarchaeota | Thaumarchaeota | *Nitrosopumilus* | | 8.3 |
| Proteobacteria | Gammaproteobacteria | *Candidatus* Endobugula | glebosa | 5.2 |
| Verrucomicrobia | Opitutae | *Candidatus* Synechomycale | tutelaris | 5.0 |
| Crenarchaeota | Thaumarchaeota | *Nitrosopumilus* | *maritimus* | 2.9 |
| Proteobacteria | Gammaproteobacteria | *Methylonatrum* | *kenyense* | 2.9 |
| Proteobacteria | Alphaproteobacteria | *Azospirillum* | | 2.9 |
| Second generation sponge_Sea | | | | |
| Proteobacteria | Alphaproteobacteria | *Hyphomicrobium* | *aestuarii* | 34.9 |
| Proteobacteria | Alphaproteobacteria | *Hyphomonas* | | 28.3 |
| Verrucomicrobia | Opitutae | *Candidatus* Synechomycale | tutelaris | 5.5 |
| Crenarchaeota | Thaumarchaeota | *Nitrosopumilus* | | 5.4 |
| Proteobacteria | Gammaproteobacteria | *Candidatus* Endobugula | glebosa | 4.2 |
| Proteobacteria | Alphaproteobacteria | *Azospirillum* | | 3.0 |
| Crenarchaeota | Thaumarchaeota | *Nitrosopumilus* | *maritimus* | 1.9 |
| Proteobacteria | Gammaproteobacteria | *Methylonatrum* | *kenyense* | 0.5 |

These data were obtained in a different year than those in Fig. 4D.

**Table S8. PCR Primers used for the detection of the *myl* cluster.**

| Position | Primer Name | Sequence |
|---|---|---|
| *N*-terminus side | *myl*-Nterm-F | TCATGGGCGGTTATGAACGG |
| | *myl*-N-R | GGGTATCTTGCCCAAAGGTC |
| middle | *myl*-Mid-F | CAGATGGAACGCAGCAGG |
| | *myl*-Mid-R | GGGCTTTAAGTCGCTCCCAT |
| *C*-terminus side | *myl*-Cterm-F | CTGATGTCATGCATGGGC |
| | *myl*-Cterm-R | GCCTTGTTGTGTCATGCC |

**Table S9. Probes for fluorescence in situ hybridization analyses.**

| Probe Name | Sequence | Description | Reference |
|---|---|---|---|
| EUB388 I+ | GCTGCCTCCCGTAGGAGT | most bacteria | Daims *et al.* (1999) |
| EUB388 III+ | GCTGCCACCCGTAGGTGT | EUB388 for Verrucomicrobia | Daims *et al.* (1999) |
| Non-EUB388 I+ | ACTCCTACGGGAGGCAGC | anti-sense of EUB388 III+ | this study |
| St-16S1 | CTTACGCACTTCGGGTGG | specific for *Ca*. S. tutelaris 16S rRNA | this study |
| St-16S2 | CACGTTTGCTAAGGCCCT | specific for *Ca*. S. tutelaris 16S rRNA | this study |
| St-16S3 | CCTGACACACTTAGCGAC | specific for *Ca*. S. tutelaris 16S rRNA | this study |
| St-16S4 | ACTCCTTCCTAAGCGTCA | specific for *Ca*. S. tutelaris 16S rRNA | this study |
| St-16S5 | CCTCAGGCGGCACACTTA | specific for *Ca*. S. tutelaris 16S rRNA | this study |
